# Genetic analysis of *Aegilops tauschii*-derived seedling resistance to leaf rust in synthetic hexaploid wheat

**DOI:** 10.1101/721415

**Authors:** Volker Mohler, Michael Schmolke, Friedrich J. Zeller, Sai L.K. Hsam

## Abstract

Seedling resistance to leaf rust available in the synthetic hexaploid wheat line Syn137 was characterized by means of cytogenetic and linkage mapping. Monosomic analysis located a single dominant gene for leaf rust resistance on chromosome 5D. Molecular mapping not only confirmed this location but also positioned the gene to the distal part of the long arm of chromosome 5D. A test of allelism showed that the gene, tentatively named *LrSyn137*, is independent but closely linked to *Lr1*. It appears that Syn137 is occasionally heterogeneous for *Lr1* since the analysis of the *Lr1*-specific marker RGA567-5 in the mapping population indicated the presence of *Lr1*. Syn137 represents another source of genetic variation that can be useful for the diversification of leaf rust resistance in wheat cultivars.

## Introduction

Leaf rust, caused by the fungus *Puccinia triticina* (*Pt*), is a foliar wheat disease of global significance. The most effective, economical, and environmentally sound means of controlling this disease is the deployment of resistant wheat cultivars. To date, formally designated leaf rust resistance genes have been catalogued at 76 loci (*Lr1* – *Lr79*). The release of cultivars with resistance based upon single major resistance genes leads to the emergence of pathotypes with matching genes for virulence or that have lost genes for avirulence. Hence, the identification of new leaf rust resistant sources becomes an ongoing process to maintain resistance diversity in released cultivars. Various strategies for prolonging resistance in commercial cultivars have been proposed. These include the deployment of different combinations of major and/or adult plant resistance genes within single cultivars, referred to as gene pyramiding or gene stacking, or between different plants within the wheat crop such as in agronomically similar cultivar mixtures or genetically related multiline varieties (Burdon et al. 2014).

Synthetic hexaploid wheat lines (2n = 6x = 42, AABBDD sub-genomes) produced as chromosomally doubled hybrids (via colchicine treatment) between *Triticum turgidum* (2n = 4x = 28, AABB sub-genomes) and *Aegilops tauschii* (2n = 2x = 14, DD genome) are important genetic resources enabling the direct exploitation of the genetic variation present in both the AABB sub-genome progenitors and close relatives and the D genome progenitor of cultivated wheat. Of the currently named leaf rust resistance genes in wheat, five were derived from *Ae. tauschii*. The cloned leaf rust resistance gene *Lr21* (Huang et al. 2003) was first made available in a synthetic line RL5406 (Rowland and Kerber 1974; McIntosh et al. 1995) before being backcrossed in cultivar Thatcher and other genotypes. Introgression of genes *Lr22a* (Rowland and Kerber 1974) and *Lr32* (Kerber 1987) was achieved in a similar way, whereas *Lr39* (Raupp et al. 2001) and *Lr42* (Cox et al. 1993) were transferred by direct hybridisation with common wheat and embryo rescue from the F_1_ hybrids (Gill and Raupp 1987). Previously designated genes *Lr40* and *Lr41* were shown to be *Lr21* and *Lr39*, respectively, whereas wheat stock WGRC16 reported to have *Lr43* carried gene combination *Lr21* and *Lr39* (Gill et al. 2008). Therefore, these gene designations were deleted from the Catalogue of Gene Symbols for Wheat (McIntosh et al. 2013).

The present study addressed the genetic analysis of seedling resistance to leaf rust in the synthetic hexaploid wheat line Syn137 and involved the teamwork of traditional and molecular techniques.

## Materials and Methods

### Plant materials and genetic analysis

Syn137 (68.111/RGB-U//Ward/3/*Ae. tauschii* (WX629)), a leaf rust resistant entry in the CIMMYT 1^st^ AB×D Elite Synthetics Programme, was crossed with each of the 21 Chinese Spring (CS) monosomic lines originally developed by, and obtained from, E.R. Sears, University of Missouri, USA. Cytologically confirmed monosomic F_1_ plants were grown in the greenhouse to obtain F_2_ seeds. The location of genes by monosomic analysis depends on the identification of an abnormal genetic ratio in one cross (the ‘critical’ cross) in which the resistance gene is located on the non-pairing monosomic chromosome, compared with normal disomic inheritance of the resistance gene in the 20 ‘non-critical’ crosses. To confirm F_2_ segregation of the critical cross, F_2:3_ lines (34 plants each) of 20 resistant F_2_ plants were assessed for response to leaf rust. A total of 93 F_2:3_ lines originating from the disomic cross between CS and Syn137 were used for analysis of linkage of molecular markers and the resistance gene. Chi-squared tests for goodness of fit were used to test for deviation of observed data from theoretically expected segregation ratios. Chi-square values were corrected for continuity (http://vassarstats.net/csfit.html). A test of allelism between the gene in Syn137 and *Lr1* in the Thatcher derivative RL 6003 involved 54 F_2:3_ lines (24 to 30 seedlings each) and deployed rust isolates S12 (avirulent to both resistance genes) and Pt60 (virulent to *Lr1* and avirulent to the gene in Syn137). A chi-square test of independence using a genetic ratio of 11 (homozygous resistant + segregating 15: 1): 2 (segregating 3: 1, resistant to both isolates): 2 (segregating 3: 1 to one isolate, homozygous susceptible to the other): 1 (homozygous susceptible) was applied to assess linkage of the two resistance genes. A set of 13 single-gene lines with known leaf rust resistance genes was used to compare leaf rust responses to Syn137.

### Leaf rust reaction tests

Disease testing was carried out on primary leaves of 10-days old host seedlings according to the method of Felsenstein et al. (1998). The 3 cm-leaf segments were cultured in clear polystyrene boxes on 6g/l agar and 35mg/l benzimidazole. The experiments used single spore progenies, mostly collected in Europe. *Pt* isolate Race 9 was originally provided by P.L. Dyck, Winnipeg, Canada. Inoculum was produced on the susceptible wheat cultivar Kanzler, collected, and dispersed above the exposed leaves in a settling tower at densities of 400-500 spores/cm^2^. Plates with inoculated leaf segments were wrapped in paper towel wetted with distilled water, and then enclosed in black plastic for 24 h. The leaf segment boxes were maintained under continuous light in a growth chamber at 17°C and at 60–80% relative humidity. Disease response was measured 10 days after inoculation, and followed the 0–4 infection type (IT) scoring system, in which IT ‘0’ indicated no visible symptoms, IT ‘;’ indicated hypersensitive flecks, IT ‘1’ indicated small uredinia with necrosis, IT ‘2’ indicated small to medium sized uredinia with green islands and surrounded by necrosis or chlorosis, IT ‘3’ indicated medium to large sized uredinia with chlorosis, IT ‘4’ indicated large uredinia without chlorosis, and IT ‘X’ indicated heterogeneous ITs, similarly distributed over a given leaf. Plus and minus signs were used to indicate higher and lower response than average for a given IT. Infection types ‘3’ or higher were regarded as compatible (high IT), whereas ITs of ‘2’ or lower were regarded as incompatible (low IT).

### Molecular mapping

Bulked segregant analysis (Michelmore et al. 1991) was used to identify microsatellite marker loci from wheat chromosome 5D with linkage to the gene in Syn137. Resistant and susceptible bulks consisted of DNA from eight homozygous resistant and eight homozygous susceptible F_2:3_ lines of CS × Syn137 mapping population. Fifteen individuals of each F_2:3_ line were collected for DNA extraction (Huang et al. 2000b). Analysis of microsatellite markers from chromosome 5D was carried out as described in Huang et al. (2000a). Primer information and PCR conditions for the *Lr1*-specific marker RGA567-5 were taken from Cloutier et al. (2008). Following segregation analysis of leaf rust response and molecular markers in the mapping population, a partial linkage map was computed with the program JoinMap 5.0. Map distances were calculated using the Haldane function. Charts of genetic linkage maps were drawn with the computer program MapChart 2.1 (Voorrips 2002).

## Results

### Leaf rust response

The response to 9 *Pt* isolates of 13 wheat cultivars and lines with documented resistance genes was compared to that of Syn137 employing agar-mounted primary leaf segment tests. The IT pattern of Syn137 was different to those obtained for the 13 reference genotypes (Table 1). The resistance gene in Syn137 was characterized by low ITs to all *Pt* cultures ranging from ‘0;’ to ‘;2=’.

**Table 1:** Leaf rust infection types produced by wheat cultivars/lines possessing known leaf rust resistance genes and Syn137 after inoculation with nine *Pt* isolates

| Line | *Pt* isolate |  |  |  |  |  |  |  |  | *Lr* gene |
| --- | --- | --- | --- | --- | --- | --- | --- | --- | --- | --- |
|  | S12 | S28 | S29 | S48 | S71 | Pt8 | Pt9 | Pt60 | Race 9 |  |
| Syn137 | ;1 | ;1 | ;1= | 12= | ;2= | ; | ;1 | 0; | 0; | *Syn137* |
| RL 6003<sup>1</sup> | 0; | 1 | 0; | 0; | 0; | 0; | 0; | 3++ | 0; | 1 |
| RL 6016<sup>1</sup> | ; | 12 | ; | ; | ; | 1 | 1 | 3++ | 2 | 2a |
| Democrat | 2 | 2+ | 3 | 3 | 1 | X+ | 3 | ; | 12 | 3a |
| Klein Aniversario | 2 | 2 | 3 | 3= | 3 | ; | 3 | 1 | 3 | 3c |
| RL 6010<sup>1</sup> | 0; | 2 | 0; | 0; | 0; | 0; | 0; | 3= | 0; | 9 |
| Kenya 1-12E-19J | 4 | ;1 | 3 | 2 | 3 | X | 3 | 4 | 4 | 15 |
| Exchange | 4 | 4 | 2 | 3 | X | 2 | 3 | X | 4 | 16 |
| Agatha | 0; | 0; | 0; | 0; | 0; | 0; | 0; | 0; | 0; | 19 |
| Thew | 3 | 2 | 0 | 3 | 3 | 2 | 3 | 4 | 3 | 20 |
| RL 5289<sup>1</sup> | 3 | 2 | 3 | 3 | 1 | 3++ | 3 | 12 | 3 | 21 |
| Agent | ; | ;1 | ; | ; | ; | 1 | 1 | 1 | 1 | 24 |
| Disponent | 3++ | 1 | ; | 2 | 0; | 3 | 4 | 12 | 2 | 26 |
| RL 6049<sup>1</sup> | 3 | 1 | 1 | 3 | 3 | 2 | 2 | ;1 | 2 | 30 |

<sup>1</sup> Registered accessions of Agriculture Canada Research Station, Winnipeg

### Monosomic analysis

F_2_ populations from the monsomic F_1_ hybrids were tested with *Pt* isolate S12. Segregation in all crosses, except that involving chromosome 5D, corresponded to that expected for 3 resistant: 1 susceptible, indicating a single dominant gene for resistance (Table 2). Segregation in the 5D cross deviated significantly from 3: 1 (χ^2^_3:1_ = 21.49, *P* < 0.0001, df = 1), with only 1 of 76 seedlings scored as susceptible. In this critical cross it was expected that the disomics (RR) and monosomics (R-) were resistant, whereas the nullisomics (--) susceptible. Hence, it is assumed that the susceptible plant was a nullisomic indicating that the resistance gene was located on chromosome 5D. In addition, 20 F_3_ lines derived from randomly selected resistant F_2_ plants of the critical cross were progeny tested. It has been well documented that the accuracy of individual F_2_ plant classification can be established on the basis of progeny testing. Where a single chromosome conferring resistance is involved, F_2_ progenies of non-critical crosses should be segregating 1 resistant: 2 segregating: 1 susceptible, while the progeny of a critical cross should have a reduced number of the latter (McIntosh 1987). In the present result, none of the F_2_ progeny showed a 1: 2: 1 segregation confirming that the resistance gene is located on chromosome 5D.

**Table 2:** F_2_ segregation for seedling reaction to *Pt* isolate S12 in progenies of monosomic F_1_ plants from crosses between Chinese Spring monosomics and Syn137

| Monosomic cross | Observed segregation | | $\chi^2_{3:1}$ | *P* |
| --- | --- | --- | --- | --- |
|  | Resistant | Susceptible |  |  |
| 1A × Syn137 | 64 | 15 | 1.21 | 0.2713 |
| 2A × Syn137 | 37 | 11 | 0.03 | 0.8625 |
| 3A × Syn137 | 63 | 19 | 0.07 | 0.7913 |
| 4A × Syn137 | 70 | 21 | 0.09 | 0.7642 |
| 5A × Syn137 | 67 | 25 | 0.13 | 0.7184 |
| 6A × Syn137 | 70 | 19 | 0.45 | 0.5023 |
| 7A × Syn137 | 71 | 17 | 1.23 | 0.2674 |
| 1B × Syn137 | 58 | 24 | 0.59 | 0.4424 |
| 2B × Syn137 | 59 | 15 | 0.65 | 0.4201 |
| 3B × Syn137 | 55 | 25 | 1.35 | 0.2453 |
| 4B × Syn137 | 77 | 17 | 2.04 | 0.1532 |
| 5B × Syn137 | 68 | 20 | 0.13 | 0.7184 |
| 6B × Syn137 | 65 | 20 | 0.04 | 0.8415 |
| 7B × Syn137 | 61 | 24 | 0.32 | 0.5716 |
| 1D × Syn137 | 73 | 18 | 1.05 | 0.3055 |
| 2D × Syn137 | 69 | 23 | 0.01 | 1.0000 |
| 3D × Syn137 | 63 | 24 | 0.19 | 0.6629 |
| 4D × Syn137 | 60 | 18 | 0.07 | 0.7913 |
| 5D × Syn137 | 75 | 1 | 21.49 | <.0001 |
| 6D × Syn137 | 63 | 18 | 0.20 | 0.6547 |
| 7D × Syn137 | 67 | 22 | 0.00 | 1.0000 |
| Total excluding 5D | 1280 | 395 | 1.72 | 0.1897 |

### Genetic mapping

Assuming single-gene segregation, the F_3_ population of cross CS × Syn137 displayed distorted segregation when tested with isolates S12 and Race 9 (26 homozygous resistant: 61 heterozygous: 6 susceptible; χ^2^_1:2:1_ = 17.65, *P* = 0.0001, df = 2). Based on the results of monosomic analysis, microsatellite markers evenly distributed across chromosome 5D were used for molecular analysis. The five microsatellite marker loci *Xbarc177*, *Xgwm269*, *Xgwm272*, *Xgwm565* and *Xgwm654* from the long arm of chromosome 5D were polymorphic in bulked segregant analysis. Linkage analysis of phenotypic and molecular data in 93 F_2:3_ lines from CS × Syn137 refined location of the resistance gene distal to *Xgwm272* (Fig. 1). As the dominant leaf rust resistance gene *Lr1* is also known to be located on chromosome 5DL, marker RGA567-5 functional for *Lr1* was assayed on the parental lines. Syn137 showed amplification of the RGA567-5 marker fragment, whereas CS was null. Segregation analysis across the population showed that the *Lr1*-specific marker mapped 5.6 cM proximal to the studied resistance locus (Fig. 1). The new leaf rust resistance gene was temporarily designated *LrSyn137*. As observed for the studied phenotype, all marker loci deviated significantly from Mendelian expectations (Table 3). While all loci showed a deficiency of CS – the female parent – homozygotes, *LrSyn137* showed an excess of heterozygotes and the molecular marker loci were skewed towards homozygous Syn137 genotypes.

**Fig. 1:**
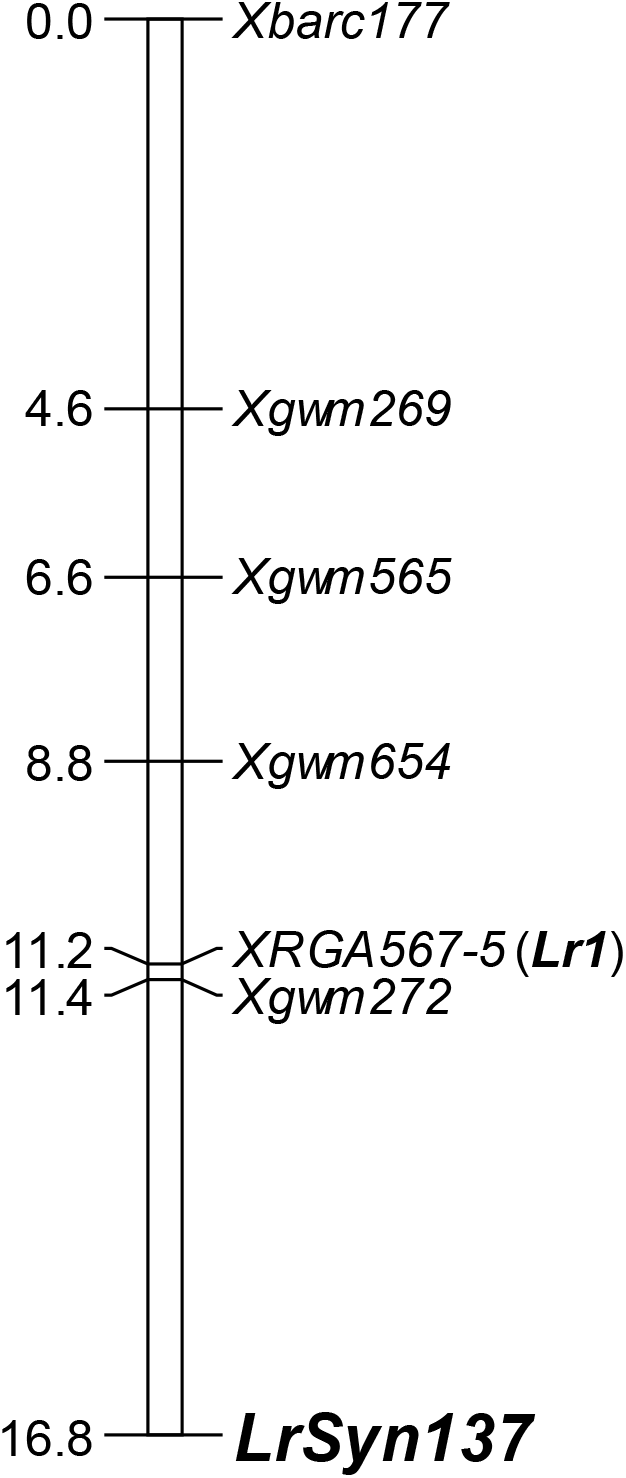
Linkage map of the distal part of wheat chromosome 5DL including leaf rust resistance gene *LrSyn137*. Absolute map positions in cM and marker names are shown on the left and right sides, respectively, of the genetic map.

**Table 3:**
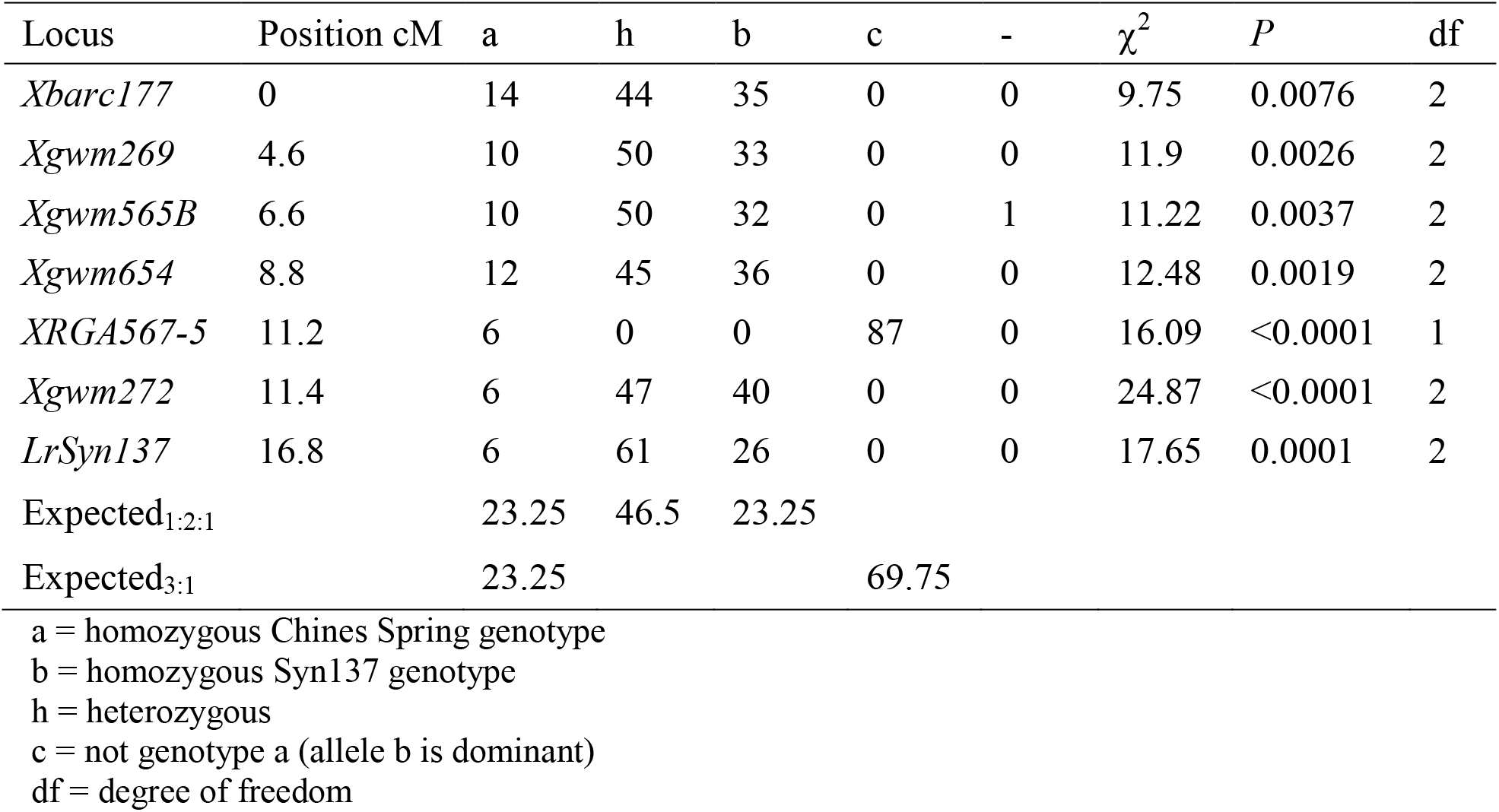
Locus genotype frequencies in CS × Syn137 population

### Test of allelism

A test of allelism was conducted between *LrSyn137* and *Lr1*. Among the 54 F_3_ progeny tested with *Pt* isolates S12, to which both genes showed resistance, and Pt60, which was virulent to *Lr1* but avirulent to *LrSyn137* (Table 1), the following segregation was obtained: 18 families were either homozygous resistant or segregating 15 resistant: 1 susceptible; 25 families showed a 3: 1, resistant to susceptible, segregation pattern to *Pt* isolate S12, to which both wheat lines showed resistance; 10 families segregated into 3 resistant: 1 susceptible to *Pt* isolate S12, but these families were concurrently susceptible to *Pt* isolate Pt60 and one family was homozygous susceptible to both *Pt* isolates. A chi-square test of independence of the pooled data using a genetic ratio of 11: 2: 2: 1 indicated that the two resistance loci were linked (χ^2^_11:2:2:1_ = 62.43, *P* < 0.0001, df = 3). Moreover, when considering only *Pt* isolate Pt60, a segregation ratio of 3: 1 (42 resistant and 12 susceptible F_3_ families; χ^2^_3:1_ = 0.22, *P* = 0.76, df = 1) was obtained. This result clearly showed that *LrSyn137* is inherited in a dominant manner.

## Discussion

Cytogenetic and linkage mapping located a dominant leaf rust resistance gene in Syn137 on chromosome 5D. In the course of determining the identity of the resistance gene on chromosome 5DL, a functional marker for *Lr1* was given priority to be assayed on the mapping population. The *Lr1*-specific marker RGA567-5 was found to map proximal to *LrSyn137* indicating distinctiveness of *LrSyn137* from *Lr1*. In addition, close linkage of RGA567-5, and thus *Lr1*, proximal to SSR marker locus *Xgwm272* could be confirmed (Ling et al. 2003). Despite a limited number of progeny, a genetic test of allelism between *Lr1* and *LrSyn137* further supported that the two genes are linked to each other.

The experiments conducted, however, suggested that Syn137 seems to be heterogeneous for resistance gene *Lr1*. However, heterogeneity for *Lr1* seems to be rare as only the line that was used for establishing the mapping population seemed to have carried *Lr1*; besides the successful allelic cross, all non-critical monosomic crosses – *Pt* isolate S12 was avirulent to both *LrSyn137* and *Lr1* – have shown single-gene segregation. The mapping population showed segregation distortion of all loci on chromosome 5DL. Therefore, it appears that compared to lines possessing only *LrSyn137*, the line carrying both *LrSyn137* and *Lr1*, additionally carries genes on chromosome 5DL generating a distortion in normal segregation in favor of themselves. Similar to our observations, Faris et al. (1998) and Li et al. (2015) reported distorter loci in *Ae. tauschii* and common wheat, respectively, located in the same genomic region on chromosome 5DL.

Three formally designated genes were located on chromosome 5D: *Lr1*, shown to be available in many wheat cultivars (McIntosh et al. 1995) and *Ae. tauschii* accessions (e.g., Ling et al. 2004), *Lr57* from *Ae. geniculata* (Kuraparthy et al. 2007), and *Lr70* (Hiebert et al. 2014) from common wheat; of which the latter two were assigned to the short arm of chromosome 5D. Qi et al. (2015) described leaf rust resistance gene *LrLB88* on chromosome 5DL that co-segregated with *Lr1* but showed a reaction pattern to 13 Chinese *Pt* pathotypes that was clearly distinct to *Lr1*. Whether *LrSyn137* and *LrLB88* are independent genes or *LrLB88* is an allele or closely linked to *Lr1* needs to be determined in follow-up research. However, satisfactory evidence is presented, similar to powdery mildew (Miranda et al. 2006, 2007), that diverse leaf rust resistance genes are located in the terminal region of chromosome 5DL. Synthetic hexaploid wheats were mainly used for the transfer of genes controlling resistance to biotic stress because of their mostly simple inheritance and ease of detection; but they have also emerged as a valuable resource for enhancing tolerance to abiotic stresses, nutritional value and grain quality attributes (van Ginkel and Ogbonnaya 2007; Li et al. 2018). However, new avenues must be taken for increasing allele diversity and recombination in structured populations to manage the wealth of information available in synthetic wheats. Therefore, population types such as the multiparent advanced generation intercross (Cavanagh et al. 2008) or multiple synthetic derivatives (Gorafi et al. 2018) will support introgression breeding and accelerate gene discovery.

